# Cerebral perfusion alterations in temporal lobe epilepsy: Structural underpinnings and network disruptions

**DOI:** 10.1101/2023.08.22.553552

**Authors:** Alexander Ngo, Jessica Royer, Raúl Rodríguez-Cruces, Ke Xie, Jordan DeKraker, Hans Auer, Shahin Tavakol, Jack Lam, Dewi Schrader, Roy W. R. Dudley, Andrea Bernasconi, Neda Bernasconi, Birgit Frauscher, Sara Larivière, Boris C. Bernhardt

## Abstract

**Objective:** Neuroimaging has been the prevailing method to study brain networks in temporal lobe epilepsy (TLE), showing widespread alterations beyond the mesiotemporal lobe. Despite the critical role of the cerebrovascular system in maintaining whole-brain structure and function, changes in cerebral blood flow (CBF) remain incompletely understood in the disease.

**Methods:** We studied 24 individuals with pharmaco-resistant TLE and 38 healthy adults using multimodal 3T magnetic resonance imaging. We compared regional CBF changes in patients relative to controls and related our perfusion findings to morphological and microstructural metrics. We further probed inter-regional vascular networks in TLE, using graph theoretical CBF covariance analysis. Finally, we assessed the effects of disease duration to study progressive changes.

**Results:** Compared to controls, individuals with TLE showed widespread CBF reductions, predominantly in fronto-temporal regions, with 83% of patients showing more marked decreases ipsilateral than contralateral to the seizure focus. Parallel structural profiling and network-based models showed that cerebral hypoperfusion may be partly constrained by grey and white matter changes and topologically segregated from whole-brain perfusion networks. Negative effects of progressive disease duration further targeted regional CBF profiles in patients. Findings were confirmed in a subgroup of patients who remained seizure-free after surgery.

**Interpretation:** Our multimodal findings provide insights into vascular contributions to TLE pathophysiology and highlight their clinical potential in seizure lateralization.

## Introduction

Temporal lobe epilepsy (TLE) is one of the most common pharmaco-resistant epilepsies in adults. Traditionally considered a “focal” disorder with a mesiotemporal epicentre, culminating multimodal neuroimaging research has demonstrated widespread alterations across allocortical, neocortical, and subcortical territories,^1-4^ playing an integral part in conceptualizing TLE as a disease of large-scale networks.^5^

Cerebral blood flow (CBF) indicates blood perfusion and is, thus, key for healthy brain structure and network organization.^6, 7^ Arterial spin labelling (ASL) magnetically labels water proton spins of the inflowing arterial blood and offers a non-invasive window into CBF.^8^ In epilepsy, perfusion alterations are modulated by ictogenesis, often showing initial hyperperfusion, followed by post- and inter-ictal hypoperfusion in the epileptogenic focus.^9-11^ Mapping these changes in TLE interictally, in particular, has identified decreased and asymmetric CBF, often spanning most of the affected temporal region.^9, 11-18^ Despite being characterized as a network disorder, prior ASL studies have mainly focused on the temporal lobe using volume-based approaches. Brain-wide surface-based analysis, in contrast, provides an unmatched avenue to map CBF profiles at a continuous and region-specific level, with advanced capabilities for cross-subject alignment. Precision mapping of perfusion changes in and beyond the seizure focus can help shed light on large-scale pathophysiological effects of TLE, while providing a surface-based reference frame to assess the interplay between altered vascular supply and underlying structural brain properties.

Controversy exists regarding the mechanisms contributing to inter-ictal hypoperfusion in TLE. While alterations are generally hypothesized to reflect atypical microstructural and morphology commonly seen in patients,^19^ prior studies have reported a dissociation between structural damage and CBF.^11, 13, 14^ Surface-based analysis of CBF, therefore, presents a systematic and quantitative gateway to run multimodal integrative studies that link disruptions in blood flow with morphological and microstructural damage—core features of the disease—and ultimately to broaden our understanding of the pathological mechanisms underlying inter-ictal hypoperfusion.

While regional analysis of perfusion interrogates local changes in brain physiology, inter-regional associations can assess the coupling between circuit function and arterial supply.^20^ In particular, correlation analysis of CBF across participants has been associated with functional networks and may capture physiological contributions.^7, 21^ Combined with graph theoretical methods, this approach can quantify topological and organizational properties of healthy and pathological inter-connected perfusion networks.^7^ Applied to TLE, prior work has revealed decreased global efficiency and disconnections of the mesiotemporal lobe from the remaining network.^22^ Despite evidence for topological changes in perfusion networks, studying large-scale vascular reorganization, however, remains in its infancy. Complementing regional perfusion profiles with information on network topology can help identify macroscale alterations in vascular supply in TLE.

Our study mapped CBF changes in TLE relative to healthy individuals based on whole-brain ASL. To explore potential structural underpinnings of vascular compromise, we integrated surface-based CBF patterns with MRI-derived measures of grey matter morphology and superficial white matter microstructure. We further generated large-scale vascular networks based on CBF covariance analyses and quantified their topological reorganization using graph theory. To examine progressive changes, we examined disease duration effects on regional and inter-regional perfusion patterns.

## Material And Methods

### Participants

We studied 24 adult patients with pharmaco-resistant unilateral TLE (12 males, mean ± standard deviation [SD] age = 35.8 ± 10.6 years, 14 left-sided focus) and 38 age- and sex-matched healthy controls with no history of neurological or psychiatric conditions (18 males, mean ± SD age = 34.8 ± 9.3 years). All individuals agreed to participate in our imaging studies, with parameters similar to the MICA-MICs dataset,^23^ and were consecutively selected based on clinical inclusion criteria and availability of ASL scans. Participants were recruited between 2018 and 2023. Detailed clinical information is presented in **Table 1**. TLE diagnosis was determined according to the classification of the International League Against Epilepsy, based on a comprehensive examination including detailed clinical history, neurological examination, review of medical records, prolonged video-electroencephalographic (EEG) telemetry recordings, clinical neuroimaging, and neuropsychological evaluation. In summary, patients had a mean ± SD age of seizure onset of 20.2 ± 12.0 years (range = 0.5 – 49 years) and a mean ± SD epilepsy duration of 15.6 ± 11.6 years (range = 1 – 45 years). Seven participants (29.2%) had a history of childhood febrile convulsion and two (8.3%) had mild head trauma. At the time of the study, eleven patients (46.8%) had undergone neurosurgical intervention (two had undergone a selective amygdalo-hippocampectomy, six had undergone cortico-amygdalo-hippocampectomy, one had undergone surgical resection of ganglioglioma within the mesiotemporal lobe, and two had undergone stereotactic radiofrequency thermocoagulation within the mesiotemporal lobe during stereo-electroencephalography investigation). The remaining patients are surgical candidates currently awaiting treatment. MRI data were acquired prior to surgery in all patients. Since surgery with a mean ± SD follow-up time of 20.2 ± 18.5 months, eight patients (72.7%) have been completely seizure-free (Engel-IA), one (9.1%) has rare disabling seizures (Engel-IIB), one (9.1%) has no worthwhile improvements (Engel-IV), and one (9.1%) was lost for follow up. Based on established histopathological criteria, six of the nine available specimens (66.7%) showed mesiotemporal sclerosis or gliosis.

**Table 1.** Patient-specific demographic and clinical information.

| Patient | Age | Sex | TLE<br>laterality | Age<br>of<br>onset | Disease<br>duration | MRI diagnosis | Surgery | Histopathology | Engel<br>outcome |
| --- | --- | --- | --- | --- | --- | --- | --- | --- | --- |
| 1 | 33 | F | L | 27 | 6 | MTS | — | — | — |
| 2 | 41 | F | R | 16 | 25 | MTS | CAH | MTS | 1a |
| 3 | 31 | M | L | 17 | 14 | Suspected FCD in<br>parahippocampal gyrus | — | — | — |
| 4 | 52 | M | L | 49 | 3 | T2/FLAIR hyperintensity in<br>hippocampus and amygdala | — | — | — |
| 5 | 55 | F | L | 42 | 13 | T2/FLAIR hyperintensity in<br>hippocampus | SAH | MTS | 1a |
| 6 | 25 | M | R | 2 | 23 | T2/FLAIR hyperintensity in<br>hippocampus and amygdala | — | — | — |
| 7 | 53 | F | L | 8 | 45 | MTS | — | — | — |
| 8 | 20 | F | L | 15 | 5 | Ganglioglioma in uncus | Tumour<br>resection | Ganglioglioma | 1a |
| 9 | 37 | F | R | N/A | N/A | MTS | — | — | — |
| 10 | 43 | M | R | 30 | 13 | MTS | CAH | MTS | 1a |
| 11 | 35 | M | L | 22 | 13 | MRI-negative | CAH | FCD IIA | 1a |
| 12 | 32 | M | R | 18 | 14 | MRI-negative | RFTC | — | 1a |
| 13 | 41 | M | L | 14 | 27 | T2/FLAIR hyperintensity in mesiotemporal lobe and insula | RFTC | — | 4 |
| 14 | 27 | F | L | 15 | 12 | MRI-negative | — | — | — |
| 15 | 25 | F | R | 17 | 8 | FCD in fusiform gyrus and parahippocampus | CAH | Mild FCD | 2b |
| 16 | 44 | M | L | 0.5 | 43.5 | MTS | CAH | MTS | N/A |
| 17 | 28 | M | L | 27 | 1 | T2/FLAIR hyperintensity in hippocampus | — | — | — |
| 18 | 29 | M | R | 22 | 7 | MRI-negative | — | — | — |
| 19 | 27 | F | L | 13 | 14 | MRI-negative | — | — | — |
| 20 | 39 | M | R | 24 | 15 | MTS | — | — | — |
| 21 | 54 | M | L | 42 | 12 | MTS | SAH | MTS | 1a |
| 22 | 39 | F | R | 12 | 27 | MTS | CAH | MTS | 1a |
| 23 | 18 | F | L | 14 | 4 | MRI-negative | — | — | — |
| 24 | 32 | F | R | 18 | 14 | T2/FLAIR hyperintensity in<br>mesiotemporal lobe | — | — | — |

CAH = cortico-amygdalohippocampectomy; F = female; FCD = focal cortical dysplasia; L = left; M = male; MTS = mesiotemporal sclerosis; N/A = not available; R = right; RFTC = stereotactic radiofrequency thermocoagulation; SAH = selective amygdalohippocampectomy

This study was approved by the Research Ethics Board of the Montreal Neurological Institute and Hospital. All participants provided their written informed consent.

### MRI acquisition

Multimodal MRI data in patients and controls were collected on a 3T Siemens Magnetom Prisma-Fit equipped with a 64-channel head coil. Images included (*i*) high-resolution 3D T1-weighted (T1w) scans using a magnetization-prepared rapid gradient-echo (MPRAGE) sequence (repetition time [TR] = 2300 ms, echo time [TE] = 3.14 ms, flip angle = 9°, matrix = 320 × 320, voxel size = 0.8 × 0.8 × 0.8 mm^3^, field of view [FOV] = 256 × 256 mm^2^, 224 slices), (*ii*) resting-state pseudo-continuous ASL (pCASL) imaging (TR = 4150 ms, TE = 10 ms, flip angle = 90°, voxel size = 4.5 × 4.5 × 7 mm^3^, FOV = 288 × 288 mm^2^, post-label delay = 1550 ms, 14 slices) and associated magnetizations of arterial blood at equilibrium (M0; TR = 10,000 ms), and (*iii*) diffusion-weighted imaging (DWI) using a spin echo-based echo planar imaging sequence with 3 b0 images (TR = 3500 ms, TE = 64.40ms, flip angle = 90°, voxel size = 1.6×1.6×1.6 mm^3^, FOV = 224×224 mm^2^, b-values = 300, 700, and 2000 s/mm^2^, diffusion directions = 140).

### Multimodal MRI processing

Multimodal data processing was performed using the open-access micapipe^24^ (version 0.1.4; http://micapipe.readthedocs.io). Briefly, native T1-weighted (T1w) data were de-obliqued, reoriented to standard orientation (LPI: left to right, posterior to anterior, and inferior to superior), and skull stripped. Subcortical structures as well as hippocampal and neocortical surfaces were segmented from native T1w images using FSL FIRST,^25^ HippUnfold^26^ (version 1.0.0; https://hippunfold.readthedocs.io/), and Freesurfer 6.0.0^27^ (https://surfer.nmr.mgh.harvard.edu/), respectively. Subject-specific vertex-wise maps of cortical thickness were generated by measuring the Euclidean distance between corresponding pial and white matter vertices.

Diffusion MRI data were preprocessed using MRtrix3^28^ (http://www.mrtrix.org/), and included denoising, b0 intensity normalization, as well as correction for susceptibility distortion, head motion and eddy currents. As in previous works,^4, 29, 30^ we generated surfaces 2mm beneath the grey-white matter interface to target both U-fibre systems and long-range bundle terminations. Diffusion tensor-derived fractional anisotropy (FA) and mean diffusivity (MD), representative of fibre architecture and tissue microstructure, were interpolated along the surface.

ASL data were preprocessed using BASIL^31^ and FSL^25^. Structural scans were automatically cropped, bias-field corrected, skull stripped, and segmented (tissue-type and subcortical structure). We used *oxford_asl*—an automated command line tool within the BASIL toolbox—to generate calibrated maps of absolute resting-state tissue perfusion. CBF maps were co-registered to the native T1w processing space of each participant, mapped to a 10k-vertex (*i*.*e*., surface points) version of the Conte69 human symmetric surface template, and smoothed using a 10mm full-width-at-half-maximum kernel. In parallel, subject-specific subcortical and hippocampal segmentations were non-linearly registered to each individual’s native ASL space and averaged.

### Case-control perfusion analysis

Blood flow maps in individuals with TLE were *z*-scored relative to controls and sorted into ipsilateral/contralateral to the focus. As in previous work, we fitted surface-based linear models to compare brain perfusion values in patients to controls using BrainStat^32^ (version 0.4.2; https://brainstat.readthedocs.io), in MATLAB 2020b. To control for age and sex effects, the regression model for each vertex *i* was defined as:

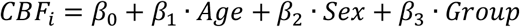

### Surface-based CBF asymmetry

To assess the clinical utility of perfusion for seizure focus lateralization, we computed interhemispheric asymmetry of individual cerebral perfusion:

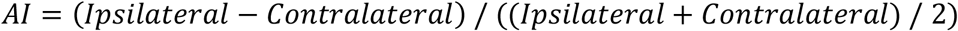

Paired *t-*tests compared ipsilateral vs contralateral CBF profiles in TLE.

### Relation to whole-brain morphology and microstructure

Surface-based linear models first compared morphometric (*i*.*e*., cortical thickness and subcortical volume) and multivariate superficial white matter microstructure measures (*i*.*e*., FA and MD) in TLE relative to controls, while controlling for age and sex. To assess blood flow alterations in TLE above and beyond the effects of grey and white matter compromise, we repeated the aforementioned perfusion analyses, while additionally correcting for thickness and microstructural alterations at each vertex. Normalized CBF values in patients were averaged before and after correction, and changes in Cohen’s *d* effect sizes were estimated.

### Graph theory analysis of vascular networks

#### i) CBF covariance networks

Subject-specific vertex-wise CBF maps were parcellated into 360 cortical areas based on the multimodal Glasser atlas,^33^ 12 subcortical regions, and bilateral hippocampi, and further corrected for age and sex. Residualized data were *z*-scored relative to controls, sorted into ipsilateral/contralateral to the focus, and used to compute covariance networks.

An interregional association matrix *R* of 374 × 374 dimensions was generated for each group (TLE and controls), where an individual entry *R*_*ij*_ (with regions *i* and *j*) contained the Pearson’s product-moment cross-correlation coefficient of CBF residuals across group-specific participants.

#### ii) Network thresholding

Prior to analysis, negative correlations and diagonal elements of *R* were set to zero. The covariance network matrices were thresholded (density range of K = 0.05 – 0.50, density interval of 0.01) to ensure an identical number of edges or wiring cost between networks in both groups^34^ and that group differences were not driven by low-level correlations.^35^

#### iii) Global network analysis

We computed two widely-used global graph theory metrics: (*i*) mean clustering coefficient (*C*), which quantifies the degree to which brain regions tend to locally interconnect with each other, and (*ii*) mean path length (*L*), which quantifies the mean shortest path (*i*.*e*., minimum number of edges) between all pairs of regions in the network.^36^ Each measure was normalized relative to the mean of 100 randomly generated networks (C_rand_ and *L*_rand_) with a similar number of nodes, edges, and degree distribution, and averaged across all cortical and subcortical regions. The area under the curve (AUC) across different sparsity thresholds was calculated to provide an integrated scalar for each network measure.

#### iv) Statistical analyses

As in previous work, network differences between patients and controls were evaluated using a nonparametric permutation test with 1,000 repetitions.^37^ In each sampling process, we randomly reassigned the perfusion data of each subject to either patient or control group, keeping the group-specific sample size unchanged, and repeated network construction and graph parameter calculations. For each measure, statistically significant differences were assessed by comparing the empirical between-group differences against the null distribution determined by the ensemble of randomized network metrics.

### Disease duration-related effects

i. To unravel the consequences of prolonged epilepsy on hypoperfusion profiles, we built linear models assessing the effects of disease duration on blood flow in TLE.
ii. Given that vascular covariance is built on group-level CBF correlations, assessing the individual effect of chronic epilepsy is challenging. We thus stratified patients by duration of illness using a median split approach (13 years, n_short_ = 9, n_long_ = 14) and reconstructed vascular networks within each subgroup. Graph theoretical properties were directly compared between short and long duration groups.

### Sensitivity analysis in seizure-free TLE

To ensure the validity of our findings, we selected patients who had undergone surgery after our initial investigation and were seizure-free at follow-up (*i*.*e*., Engel I, n = 8, mean ± SD follow-up time = 22.1 ± 20.3 months). Regional analyses of cerebral perfusion were repeated, focusing exclusively on the subgroup of post-surgical seizure-free patients.

### Multiple comparisons correction

Findings were corrected for multiple comparisons at a family-wise error (FWE) rate of *p* < 0.05 using random field theory for non-isotropic images^38^ for the cortex and at a false discovery rate^39^ (FDR) of *p* < 0.05 for subcortical structures and hippocampi.

## Results

### Blood flow reductions in TLE

In controls, we observed variations in perfusion across the brain, including higher blood flow in fronto-parieto-occipital cortices and lower levels in temporo-limbic regions (**Fig 1A**). Comparing individuals with TLE to controls revealed widespread blood flow reductions in patients, with dominant effects in ipsilateral fronto-temporal regions that extend into inferior parietal and medial occipital areas (**Fig 1B**; *p*_FWE_ < 0.05; mean ± SD Cohen’s *d* = -0.69 ± 0.15).

**Fig 1.**
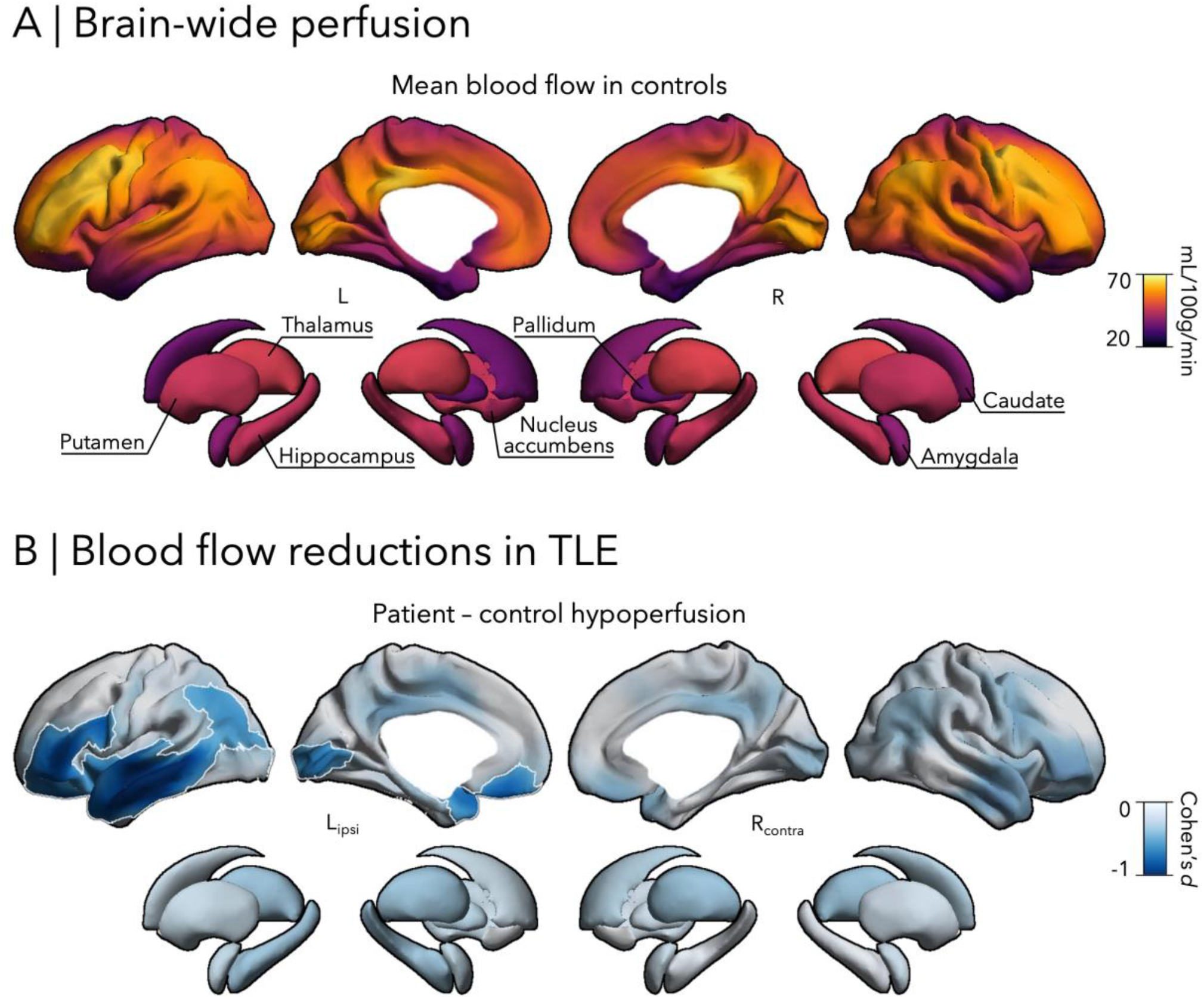
Cerebral perfusion changes in temporal lobe epilepsy (TLE). **(A)** Cortex and subcortex-wide variations in cerebral blood flow (CBF) in healthy controls (HC). **(B)** CBF alterations in patients with TLE compared to HC. Significant regions (*p*_FWE_ < 0.05) are highlighted in white.

Analyses of within-patient asymmetry maps revealed trending hypoperfusion in ipsilateral fronto-temporal regions and hippocampus (**Fig 2A**; *p*_uncorrected_ < 0.05). In particular, greater ipsilateral, compared to contralateral, reductions in the cluster were observed in a large portion of patients (**Fig 2B**; 20/24, 83.3%).

**Fig 2.**
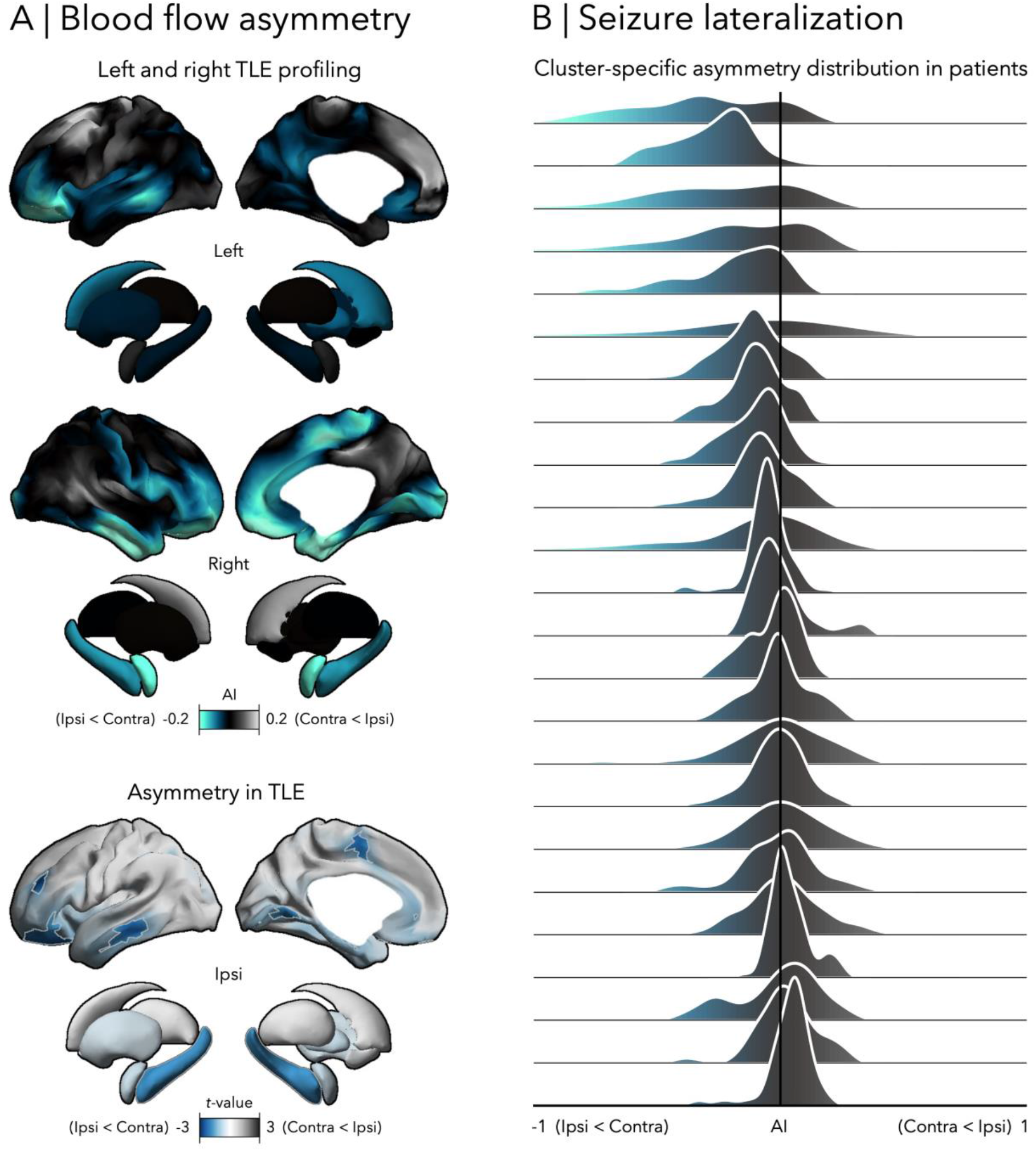
Asymmetry-based lateralization in TLE. **(A)** Perfusion asymmetry comparing ipsilateral (ipsi) versus contralateral (contra) hemispheres in left and right lateralized TLE. Trending areas (*p*_uncorrected_ < 0.05) are outlined in grey. **(B)** Patient-specific asymmetry distribution within the cluster. Black line indicates an asymmetry index (AI) of zero.

### Structural and microstructural associations

Morphological and microstructural alterations in individuals with TLE were in line with earlier imaging findings.^3, 4, 29, 30, 40^ In brief, patients showed pronounced grey matter atrophy in bilateral motor and dorsal prefrontal regions and ipsilateral hippocampus (**Fig 3A**; *p*_FWE/FDR_ < 0.05; mean ± SD Cohen’s *d* = -0.62 ± 0.11) as well as superficial white matter diffusion changes (characterized by reduced FA and increased MD) in ipsilateral temporal pole, hippocampus and putamen, and extending into bilateral frontal, temporal and occipital cortices (**Fig 3A**; *p*_FWE/FDR_ < 0.05; mean ± SD Hotelling’s *T* = 3.99 ± 0.99).

**Fig 3.**
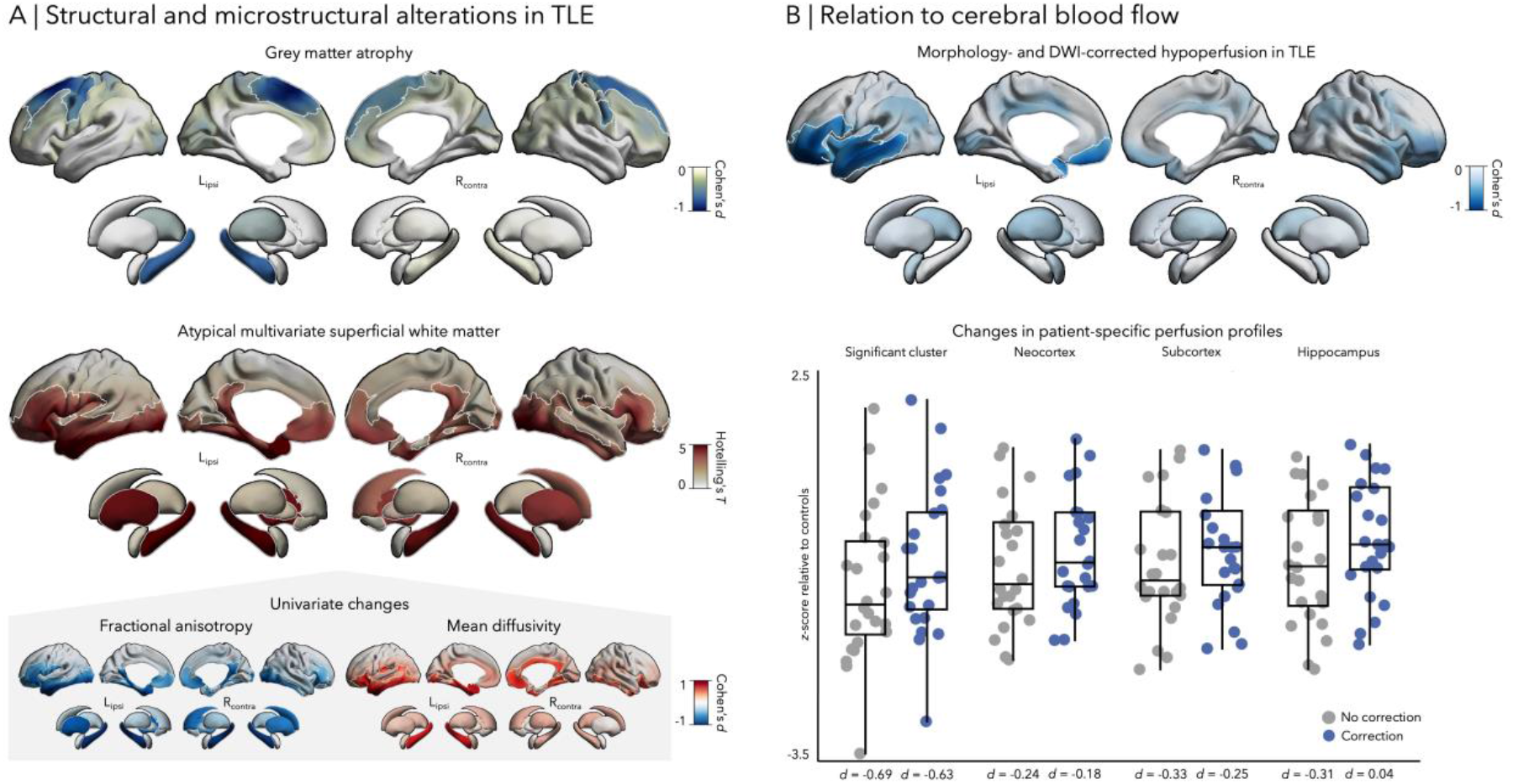
Relation to morphological and microstructural alterations. **(A)** Patterns of grey matter atrophy (derived from cortical thickness and subcortical volume) and superficial white matter changes (derived from the aggregate of FA and MD) in TLE, relative to HC. Significant differences (*p*_FWE_ < 0.05) are highlighted with white outlines. **(B)** *Top panel*: Repeating the between-group perfusion comparisons, while controlling for both morphological and microstructural alterations in each region/vertex. *Bottom panel*: Mean patient-specific profiles before (blue) and after (grey) correction are depicted for significant hypoperfusion cluster (see FFig 1B), neocortex, subcortex, and hippocampus.

While between-group comparisons of CBF patterns after correcting for MRI-derived morphological and microstructural changes revealed similar patterns compared to the main analysis (**Fig 3B**; *p*_FWE_ < 0.05), effects were overall smaller. This reduction in effect size was observed in areas of significant between-group hypoperfusion (reduction in Cohen’s *d* = -8.11%), and even more strongly across remaining subcortico-cortical territories (hippocampus = -113.64%, subcortex = -22.81%, and neocortex = -24.58%).

### Atypical vascular covariance networks in TLE

In healthy individuals, vascular networks revealed a small-world topology characterized by high clustering and short path length across the entire range of density thresholds (γ = *C/C*_rand_ > 1 and λ *= L/L*_rand_ ≈ 1) at the whole-network level (**Fig 4B**). Comparing graph theoretical metrics between patients and controls, we observed suboptimal organization in TLE, with significant decreases in clustering (*p*_FDR_ < 0.05 at *K* = 0.05–0.50; AUC = -0.17, *p* < 0.01; **Fig 4C**).

**Fig 4.**
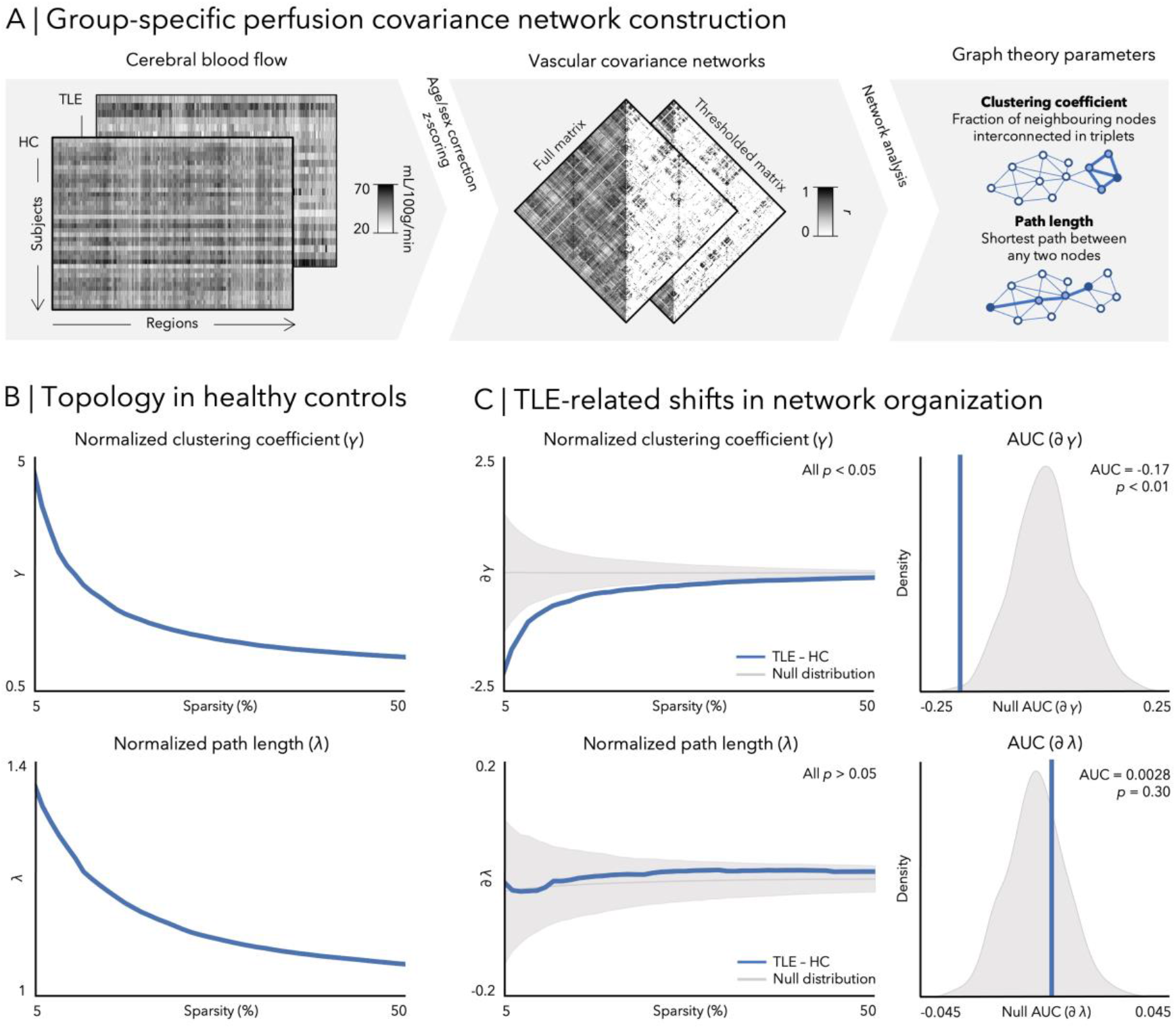
TLE changes in cerebral blood flow organization. **(A)** Schematic description of group-specific vascular covariance network construction and graph theoretical parameterization. **(B)** Global clustering coefficient (*top panel*) and path length (*bottom panel*) in healthy controls are plotted as a function of network densities (5–50%). **(C)** *Left panel*: Differences in clustering coefficient and path length between TLE and HC plotted as a function of network densities. The grey line and shaded region depict the mean and the 95% confidence interval, respectively, of the null distribution of between-group changes obtained from 1000 permutation tests at each density. Blue line indicates the empirical change in graph theory measure. *Right panel*: Area under the curve (AUC) differences across densities for clustering coefficient and path length are shown against permutation-based null differences.

### Effects of disease duration

TLE-related blood flow reductions negatively correlated with longer disease duration (whole brain *r* = -0.42, *p* = 0.05; significant cluster *r* = -0.53, *p* < 0.01). Vertex-wise associations were mainly localized in bilateral parieto-occipital cortices, ipsilateral temporal regions, hippocampus and thalamus, and contralateral orbitofrontal areas (*p*_FWE/FDR_ < 0.05; **Fig 5**).

**Fig 5.**
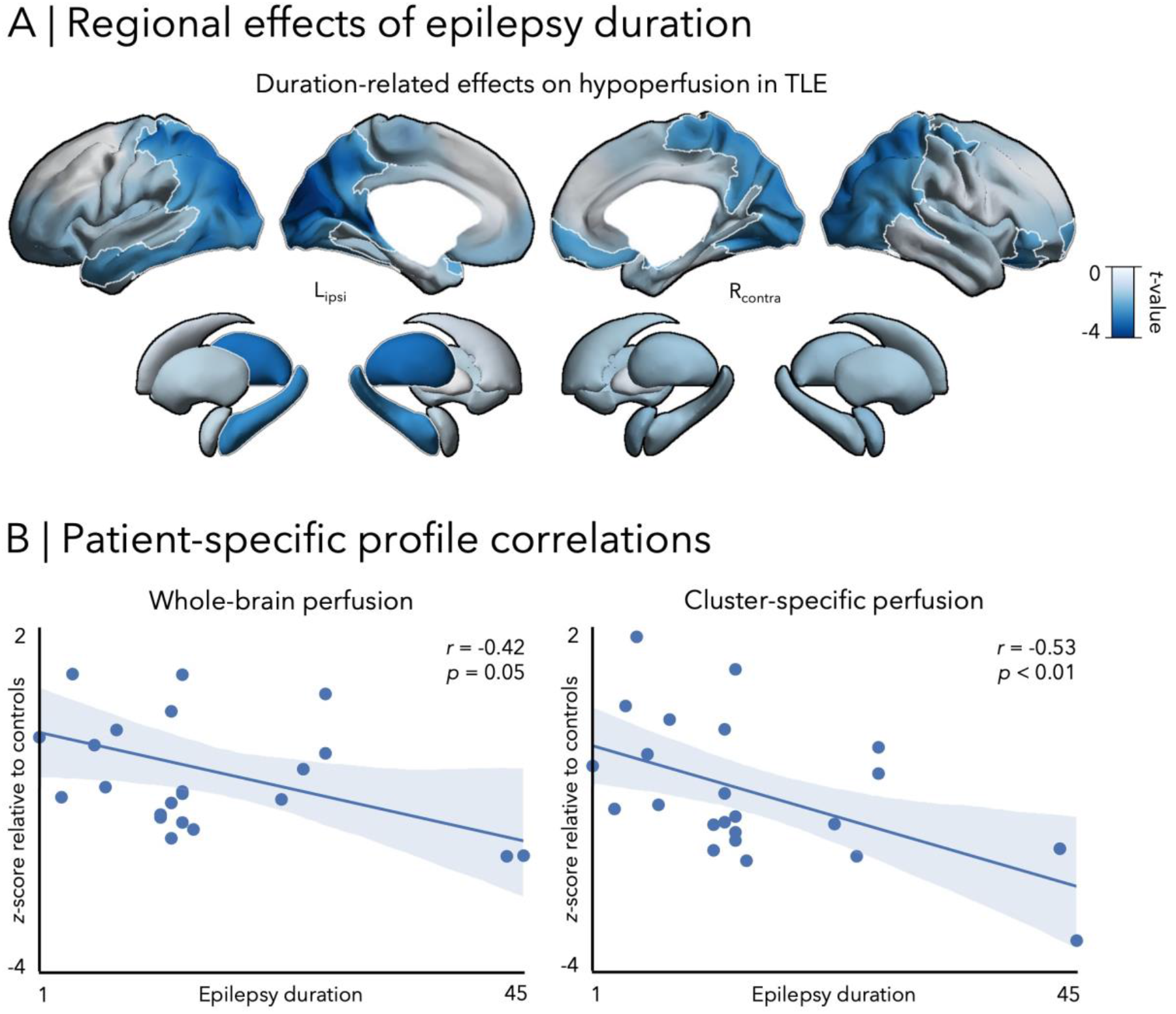
Disease duration associations with regional perfusion in TLE. **(A)** Effects of duration of epilepsy on hypoperfusion. Significant effects (*p*_FWE/FDR_ < 0.05) are outlined in white. **(B)** Correlation between duration and mean CBF across the entire brain and areas of marked hypoperfusion (see Fig 1B).

With regards to CBF covariance networks, comparison of patients grouped by short and long duration of illness revealed no significant between-group differences in clustering (*p*_uncorrected_ > 0.05 at *K* = 0.05–0.50; AUC = 0.048, *p* = 0.35) and path length (*p*_uncorrected_ > 0.05 at *K* = 0.05–0.50; AUC = 0.0096, *p* = 0.14).

### Sensitivity analysis in seizure-free patients

Repeating regional perfusion analyses in a subcohort of Engel I patients yielded virtually identical findings, showing cerebral hypoperfusion in TLE relative to controls (**Fig 6A**, *r =* 0.70, *p*_spin_ < 0.001), with strongest effects in the ipsilateral fronto-temporo-occipital regions (*p*_uncorrected_ < 0.05; mean ± SD Cohen’s *d* = -0.74 ± 0.13). Specifically, asymmetry-based lateralization showed greater ipsilateral than contralateral alterations in this cluster (**Fig 6B**; *p*_uncorrected_ < 0.05) in all seizure-free patients (8/8, 100%).

**Figure 6.**
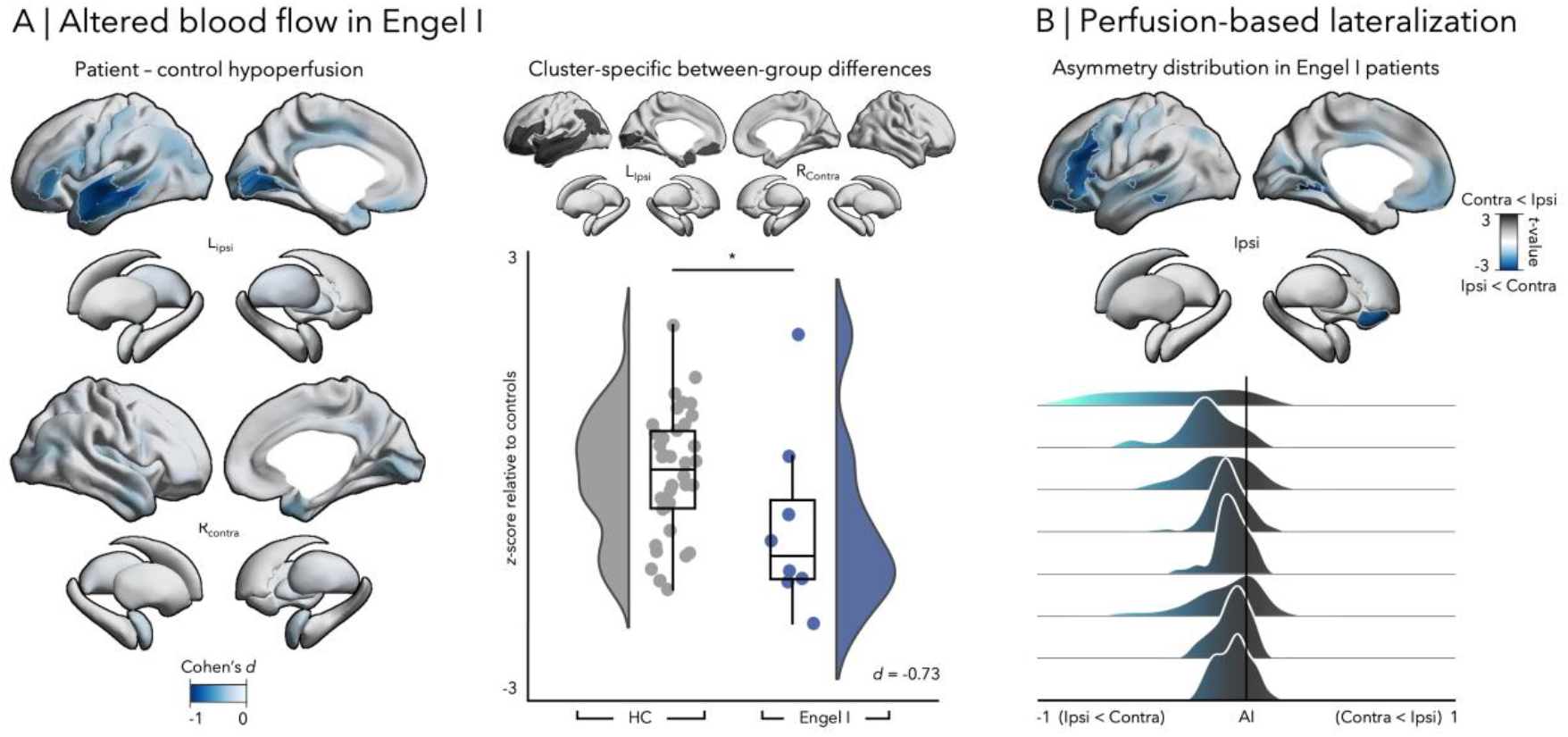
Perfusion changes in patients with post-surgical seizure freedom. **(A)** *Left panel*: CBF decreases in Engel I patients compared to HC. Trending regions (*p*_uncorrected_ < 0.05) are highlighted in grey. *Right panel*: Between-group differences in mean perfusion within the main cluster of significant hypoperfusion (see Figure 1B). Asterisk indicates a *p* < 0.05. **(B)** *Top panel*: Perfusion asymmetry maps comparing ipsi versus contra hemispheres in left and right TLE. Trending areas (*p*_uncorrected_ < 0.05) are outlined in grey. *Bottom panel*: Patient-specific asymmetry distributions within the cluster. Black line point to an AI of zero.

## Discussion

Emerging literature underscores key contributions of vasculature in TLE pathophysiology. Here, we combined regionally precise mapping of CBF with inter-regional blood flow covariance network analyses to explore large-scale perfusion imbalances. Using advanced surface-based assessment of ASL-derived tissue perfusion, together with subcortical targeting, we revealed marked decreases in whole-brain inter-ictal CBF in patients relative to controls. Patterns were spatially distributed, yet peaked in fronto-temporo-parieto-occipital regions ipsilateral to the seizure focus. At the individual level, hypoperfusion effects were localized ipsilaterally to the focus in 83% of patients, underscoring the potential clinical utility of ASL in seizure lateralization. Multimodal MRI parameter integration highlighted that hypoperfusion patterns were, in part, related to co-occurring grey matter morphological and white matter microstructural compromise. Complementing the region-specific CBF profiling, we explored vascular alterations at the macroscale with perfusion covariance analyses and graph theoretical assessment of network topology, revealing a topologically segregated network characterized by decreased clustering. Analyzing the effects of epilepsy duration further revealed predominant hypoperfusion in patients with long-standing epilepsy, possibly suggesting progressive changes in brain vasculature in the condition. Taken together, our study provides insights into vascular contributions to TLE pathophysiology and supports the clinical integration of *in vivo* perfusion mapping at regional and macroscopic scales.

While there is overall agreement on atypical perfusion in the temporal lobe ipsilateral to the seizure focus in prior literature in children and adult populations with TLE,^9, 11-18^ large-scale alterations in and extending beyond this region have not been systematically charted. Whole-brain mapping, as performed here, fills this gap by finely characterizing both temporal and extra-temporal changes in brain perfusion in TLE. We identified multi-lobar inter-ictal CBF decreases in patients relative to controls, with peak effects in ipsilateral fronto-temporal regions that extend into inferior parietal and medial occipital cortices. Such reductions may indicate reduced metabolism and atypical neurovascular coupling—a mechanism that links the metabolic demands of the brain with energy substrate supplies from the blood.^41^ Research targeting temporal regions has shown colocalized inter-ictal, ipsilateral hypoperfusion and hypometabolism in the seizure-affected area.^11, 42^ Instances of decreased blood flow have also been noted in more remote parietal, occipital, and frontal regions in some patients, possibly reflecting physiological consequences of altered neuronal activity and connectivity in chronic, inter-ictal states^4, 30^ as well as persistent reorganization secondary to seizures.^43^ Temporal and extratemporal hypoperfusion patterns have been associated with reconfigurations of tissue microvasculature. Simultaneous ictal increases in neuronal activity and metabolism may upregulate local blood flow. However, neurovascular decoupling due to seizure-induced ischemic and hypoxic attacks in post-ictal stages causes a failure to vasodilate in response to increased neuronal activity, leading to excessive constriction of vessels. This may impair vascular-scaffolding cells and flow patterns.^44^ Histological examinations of *post-mortem* specimens and surgically resected tissue in individuals with TLE and animal models point towards clear changes in microvascular morphology, including reduced vessel diameter and atypical angiogenesis.^44-46^ In conjunction with reduced metabolic demands, such alterations in vasculature of seizure-affected regions may lead to chronic hindrance of circulation and, thus, inter-ictal hypoperfusion.

Despite prior research showing consistent greater alterations ipsilateral to the seizure focus,^9, 11^-13 ASL remains clinically underutilized in relation to more common, invasive, and radioactive imaging techniques. Our analysis demonstrated asymmetry in 83% of patients, supporting the potential benefits of surface-based mapping of CBF. Notably, this may provide an accurate, inexpensive, and non-invasive tool to help resolve difficulties in lateralization of the epileptogenic zone, valuable for presurgical evaluation in patients with TLE.

Our multimodal imaging framework enabled the integration of region-specific CBF mapping with MRI-derived markers of morphology and microstructure in the same participants. In line with previous imaging studies,^3, 4, 29, 40^ TLE-related grey matter atrophy followed a bilateral and widespread cortico-subcortical topology, while microstructural alterations were primarily localized in temporo-limbic regions ipsilateral to the focus. These structural changes seem to be, in part, related to inter-ictal CBF decreases, compatible with shared pathophysiological processes. Reduction of between-group differences when taking into account structural features was most pronounced in the hippocampus, underscoring its role as the pathological epicentre of TLE.^1^ Harboring the epileptogenic zone, mesiotemporal regions are susceptible to chronic seizure activity, and as a result, display significant morphological and microstructural alterations. Acute loss of tissue and reductions in axonal density along with disruptions in the myelin sheath— energy-demanding processes dependent on blood flow—may result in inter-ictal hypometabolism. ^17, 47^ In parallel, the vascularization of the human hippocampus is unique, being the only structure that is mainly supported by two major arteries.^48^ Despite a lack of literature surrounding the effects of seizures on these large vessels, it is possible that reduced oxygen and blood demand in structurally damaged regions may cause shrinkage in these arteries. These processes may theoretically translate to blood flow decreases in the hippocampus. Conversely, extratemporal perfusion patterns in TLE may primarily arise from system-level functional alterations^4, 30^ rather than structural compromise, suggesting a different disease mechanism. As intrinsic brain function is a major determinant of regional blood flow patterns, changes in local neuronal activity and consequently altered metabolism, particularly in the neocortex, may be a core driver of hypoperfusion beyond the seizure focus. Future work combining MRI, positron emission tomography (PET), single-photon emission computerized tomography (SPECT), and ASL may help establish a clear link between brain structural, metabolic, and blood flow changes. Nonetheless, by consolidating morphological and microstructural alterations in TLE with ASL-derived perfusion mapping, our framework provides a first step towards an integrative *in vivo* perspective into multifaceted circulatory reconfigurations in the disease.

To extend regional CBF findings to the network scale, we conducted graph theoretical analyses on ASL-based perfusion correlations. Studying synchronized fluctuations in CBF may provide a novel perspective into vascular networks and their changes in TLE.^7^ We compared patients to healthy controls and demonstrated an imbalance in blood flow covariance, pointing towards topological segregation from whole-brain perfusion networks, characterized by reductions in clustering. This coupling would reflect the manifestation of a supply-and-demand principle such that blood flow is shaped by connectivity, highlighting the network-level effects of the condition. As such, this work demonstrates the potential of graph theoretical vascular covariance analytics in characterizing the network mechanisms underlying atypical blood flow organization in TLE.

Studying vascular consequences of epilepsy duration in patients, we explored progressive effects of TLE on CBF. Patterns of hypoperfusion were significantly associated with disease duration in temporo-parieto-occipital cortices, hippocampi, and thalami, with dominant effects ipsilateral to the focus. These findings align with prior cross-sectional research reporting negative correlations between perfusion alterations and longer duration of epilepsy, earlier age of onset, and higher seizure frequency.^49, 50^ Hemodynamic responses to recurrent and spreading seizures may cause vasodilation-constriction responses that erode both capillaries and small cortical arteries.^44^ Along with progressive, widespread structural and functional alterations, long-term seizure burden may lead to reduced blood flow in affected territories. Future longitudinal work tracking changes in patient-specific brain physiology, along with careful monitoring of clinical- and lifespan-related factors including medication regimen and co-morbidities, will strengthen our understanding of the vascular consequences of progressive disease burden. Collectively, mapping the effects of epilepsy severity provides new insights into the mechanisms underlying prolonged epileptogenicity and highlights the importance of early diagnosis and treatment to prevent aggravated changes in local and macroscale brain perfusion.

## Acknowledgements

A.N. acknowledges funding from the Fonds de la Recherche du Québec – Santé (FRQS) Master’s Training Scholarship. J.R. was funded by a fellowship from the Canadian Institutes of Health Research (CIHR). R.R.C. received funding from FRQS. K.X. was supported by the China Scholarship Council. J.D. was funded by a post-doctoral fellowship from the Natural Sciences and Engineering Research Council of Canada (NSERC). J.L. was supported by the McGill Faculty of Medicine Michael B. and Mary Elizabeth Wood Summer Research Bursary. R. W. R. D. received funding from FRQS, CIHR and the Foundation of McGill University Department of Neurosurgery. A.B. and N.B. were funded by FRQS and CIHR. B.F. was supported by a salary award of the FRQS (Chercheur-boursier Senior) and acknowledges support from NSERC and CIHR. S.L. was funded by a CIHR Banting Postdoctoral Fellowship and the Ann and Richard Sievers Award in Neuroscience. B.C.B. acknowledges support from NSERC, CIHR, SickKids Foundation, BrainCanada, Future Leaders Research Grant, Helmholtz International BigBrain Analytics and Learning Laboratory (HIBALL), FRQS, and the Canada Research Chairs program.

## Author Contributions

AN, SL, and BCB contributed to the conception and design of the study. AN, JR, RRC, JD, KX, HA, ST, JL, AB, NB, BF, SL, and BCB contributed to the acquisition and analysis of data. AN, RWRD, SL, and BCB contributed to drafting the text or preparing the figures.

